# APOE4, Age & Sex Regulate Respiratory Plasticity Elicited By Acute Intermittent Hypercapnic-Hypoxia

**DOI:** 10.1101/2023.01.06.522840

**Authors:** Jayakrishnan Nair, Joseph F. Welch, Alexandria B. Marciante, Tingting Hou, Qing Lu, Emily J. Fox, Gordon S. Mitchell

**Affiliations:** Breathing Research and Therapeutics Center, Department of Physical Therapy, University of Florida; Department of Biostatistics, University of Florida, Jacksonville, Florida; Brooks Rehabilitation, Jacksonville, Florida; Department of Physical Therapy, Thomas Jefferson University, PA; School of Sport, Exercise and Rehabilitation Sciences, University of Birmingham, Edgbaston, Birmingham, UK

**Keywords:** intermittent hypercapnic-hypoxia, respiratory neuroplasticity, biomarkers, genetics, *APOE_4_*, age, sex

## Abstract

**Rationale:** Acute intermittent hypoxia (AIH) is a promising strategy to induce functional motor recovery following chronic spinal cord injuries and neurodegenerative diseases. Although significant results are obtained, human AIH trials report considerable inter-individual response variability.

**Objectives:** Identify individual factors (*e.g.*, genetics, age, and sex) that determine response magnitude of healthy adults to an optimized AIH protocol, acute intermittent hypercapnic-hypoxia (AIHH).

**Methods:** Associations of individual factors with the magnitude of AIHH (15, 1-min O_2_=9.5%, CO_2_=5% episodes) induced changes in diaphragm motor-evoked potential amplitude (MEP) and inspiratory mouth occlusion pressures (P_0.1_) were evaluated in 17 healthy individuals (age=27±5 years) compared to Sham. Single nucleotide polymorphisms (SNPs) in genes linked with mechanisms of AIH induced phrenic motor plasticity (*BDNF, HTR_2A_, TPH_2_, MAOA, NTRK_2_*) and neuronal plasticity (apolipoprotein E, *APOE*) were tested. Variations in AIHH induced plasticity with age and sex were also analyzed. Additional experiments in humanized (*h*)*ApoE* knock-in rats were performed to test causality.

**Results:** AIHH-induced changes in diaphragm MEP amplitudes were lower in individuals heterozygous for *APOE_4_* (*i.e., APOE_3/4_*) allele *versus* other *APOE* genotypes (p=0.048). No significant differences were observed between any other SNPs investigated, notably *BDNFval/met* (*all p>0.05*). Males exhibited a greater diaphragm MEP enhancement *versus* females, regardless of age (p=0.004). Age was inversely related with change in P_0.1_ within the limited age range studied (p=0.007). In *hApoE_4_* knock-in rats, AIHH-induced phrenic motor plasticity was significantly lower than hApoE_3_ controls (p<0.05).

**Conclusions:** *APOE_4_* genotype, sex and age are important biological determinants of AIHH-induced respiratory motor plasticity in healthy adults.

**ADDITION TO KNOWLEDGE BASE:** Acute intermittent hypoxia (AIH) is a novel rehabilitation strategy to induce functional recovery of respiratory and non-respiratory motor systems in people with chronic spinal cord injury and/or neurodegenerative diseases. Since most AIH trials report considerable inter-individual variability in AIH outcomes, we investigated factors that potentially undermine the response to an optimized AIH protocol, acute intermittent hypercapnic-hypoxia (AIHH), in healthy humans. We demonstrate that genetics (particularly the lipid transporter, *APOE*), age and sex are important biological determinants of AIHH-induced respiratory motor plasticity.

## INTRODUCTION

Impaired breathing is a critical health concern for individuals living with lung and/or neuromuscular injury or disease. Repetitive exposures to brief episodes of low inspired O_2_ (acute intermittent hypoxia, AIH) induces respiratory motor plasticity, which can be harnessed to improve respiratory and non-respiratory motor function [1]. However, human studies published to date exhibit considerable variability in AIH responses; – 30-40% of all participants are low responders to AIH [2]. The fundamental goal of this study was to identify genetic biomarkers and the influence of age and sex on individual AIH responses in healthy humans.

In a published companion article, we reported that intermittent exposure to concurrent hypoxia and hypercapnia (AIHH: acute intermittent hypercapnic-hypoxia; ~9.5% inspired O_2_; ~4.5% inspired CO_2_) elicited robust facilitation of diaphragm motor-evoked potential, MEP, reflection volitional pathways to phrenic motor neurons, and mouth occlusion pressure in 100 msec (P_0.1_), reflecting automatic ventilatory control, in healthy adults [3]. Combined hypoxia and hypercapnia are more effective at triggering respiratory motor plasticity in humans [4, 5], possibly because greater carotid chemoreceptor activation augments serotonergic raphe neuron activity more than hypoxia alone [6, 7], and/or direct activation of raphe neurons by hypercapnia [8], thereby enhancing cell signaling cascades that strengthen synapses onto phrenic motor neurons. Consistent with published human AIH trials [2], ~40% of participants respond minimally to AIHH (defined as <25% increase in diaphragm MEP amplitudes). Since clinical trials investigating rehabilitation interventions often fail due to response heterogeneity [9–11], identifying biomarkers associated with individual responses is essential for successful large-scale clinical trials [2].

Genomic analysis has improved healthcare precision in the treatment of cancer and other clinical disorders [12]. Similar focus on identifying genetic biomarkers to align genetic profiles or individual characteristics (age or sex) with the most effective rehabilitation strategies is lacking. Genetic factors regulate AIH-induced serotonin [13] and BDNF-dependent [14] phrenic motor plasticity in rats [15, 16], leading to the hypothesis that dysfunctional genes affecting peripheral chemosensitivity, serotonergic function and/or BDNF/TrkB signaling undermine AIH-induced respiratory plasticity in humans (Figure 1). Dysfunctional genes that undermine neuroplasticity in other regions of the central nervous system, such as alleles coding for the lipid transporter apolipoprotein E (*APOE*), may also contribute to lower individual responses. For example, the *APOE_4_* isoform is associated with Alzheimer’s disease, limited recovery from neural injury, impaired glutamate receptor function and limited BDNF availability [17].

**Figure 1:**
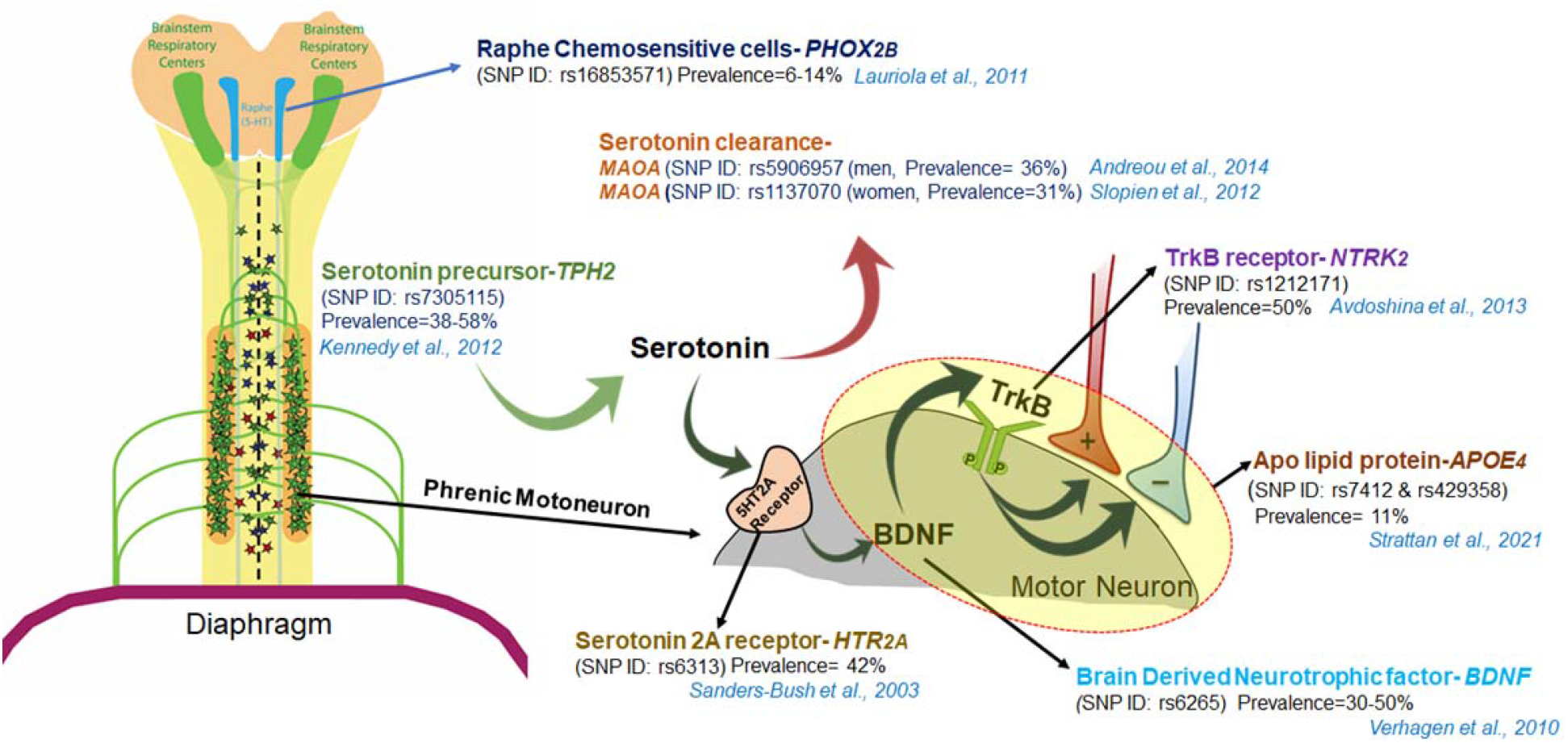
Conceptual diagram depicting cell signaling mechanisms (and candidate biomarker genes) for acute intermittent hypercapnic-hypoxia (AIHH) induced respiratory motor plasticity. The panel of SNPs with a population prevalence of >10% were tested for association with reduced AIHH-induced plasticity in humans. These include 6 SNPs in genes involved in AIH cell signaling: (1) raphe chemosensitive cells (*PHOX_2B_*), (2) serotonin precursors in the central nervous system (tryptophan hydroxylase-2, *TPH-2*), (3) serotonin clearance enzyme (monoamine oxidase A, *MAOA*), (4) serotonin-2A receptors (*HTR_2A_*), (5) brain-derived neurotrophic factor (*BDNF*) and (6) TrkB receptors (*NTRK_2_*). A seventh dysfunctional SNP in neuroplasticity related gene, APOE (*APOE_4_*), was also tested for association.

Advancing age and sex are other characteristics that differentially affect AIH-induced phrenic motor plasticity in rats [18, 19]. An age-dependent sexual dimorphism could contribute to AIH and AIHH response variability in humans. Clear links between genetics, age and sex with AIH/AIHH-induced phrenic motor plasticity in rodents informs our hypothesis that human response heterogeneity to AIHH [3] is linked with dysfunctional single nucleotide polymorphisms (SNPs) in molecules known to regulate AIH-induced phrenic motor plasticity (e.g. the *BDNFval/met* mutation) as well as age and sex.

## PROTOCOL AND METHODS

The present study was approved by the Institutional Review Board (IRB202000711) for human studies, and the Institutional Animal Care and Use Committee (IACUC202110316) for rat studies at the University of Florida. Human procedures were performed in accordance with the Declaration of Helsinki, except for registration in a database. This study is part of a larger research effort directed at optimizing AIH protocols with the use of AIHH in humans (see 3). For more information concerning methodological approaches and results, see supplementary material and Welch et al. [3].

### Participants

Seventeen participants (age range=20-40 years, mean age=27±5 years, 9 females) signed a written informed consent form to participate in the study [3]. Participants with known cardiovascular, respiratory, neurological, or infectious disease/illness, seizures, migraine (in the last 6 months), and/or metallic implants around the head, chest or shoulder region were excluded from the study. Females were screened for pregnancy. Participants were asked to refrain from caffeine consumption 8 hours prior to testing.

### Experimental Design

A detailed description of the experimental protocol and outcome measures are described elsewhere [3] and in supplementary material. Briefly, in a single-blind, cross-over sham-controlled experiment, participants received on 2 days (separated by ? 3 days): AIHH (15, 1-min hypercapnic-hypoxia episodes with 1.5 min intervals breathing room air) and normocapnic-normoxia (Sham control). During AIHH, participants inspired from a Douglas bag filled with ~9.5% O_2_ and 4.5% CO_2_ (balance N_2_). Participants breathed ambient air during Sham.

### Measures of Respiratory Neuroplasticity

Diaphragm MEPs induced by transcranial magnetic stimulation were used to assess cortico-diaphragmatic neurotransmission [3, 20, 21]. Spontaneous respiratory drive was estimated using mouth occlusion pressure in 0.1 seconds (P_0.1_) during resting breathing [22]. Tidal volume, breathing frequency and minute ventilation were also measured before (Pre), during and after (Post) AIHH and Sham. The magnitude of AIHH-induced plasticity was quantified as %-change from baseline [(Post-Pre)/Pre x100].

### Candidate Gene and Single-Nucleotide Polymorphism Selection

Based on known roles of molecules in AIH-induced phrenic motor plasticity and a minimum population penetrance of 10% [3, 23, 24], we screened for 9 SNPs in genomic DNA extracted from the subject’s saliva. Seven candidate genes (Figure 1; Table 1) included autosomal SNPs in: apolipoprotein (*APOE_4_,* SNP IDs: rs429358 [T>C] and rs7412 [T>C]), prevalence: *APOE_4_* homozygous ~11%, [17, 25, 26], *APOE_3/4_* heterozygous ~15-25% [27, 28]; brain-derived neurotrophic factor (*BDNFval/met*, SNP ID: rs6265 [C>T], prevalence ~30-50% [14, 29–31]); neurotrophic receptor tyrosine kinase 2 (*NTRK_2_*, SNP ID: rs1212171 [C>T], prevalence ~50% [32–34]); tryptophan hydroxylase 2 (*TPH_2_*, SNP ID: rs7305115, [A>G], prevalence 38-58% [35, 36]); 5-hydroxytryptamine receptor 2A (*HTR_2A_*, SNP ID: rs6313 [A>G], prevalence ~42% [37]); and, paired-like homeobox 2B (*PHOX_2B_*, SNP ID: rs16853571 [A>C], prevalence ~6-14% [38]).

**Table 1.**
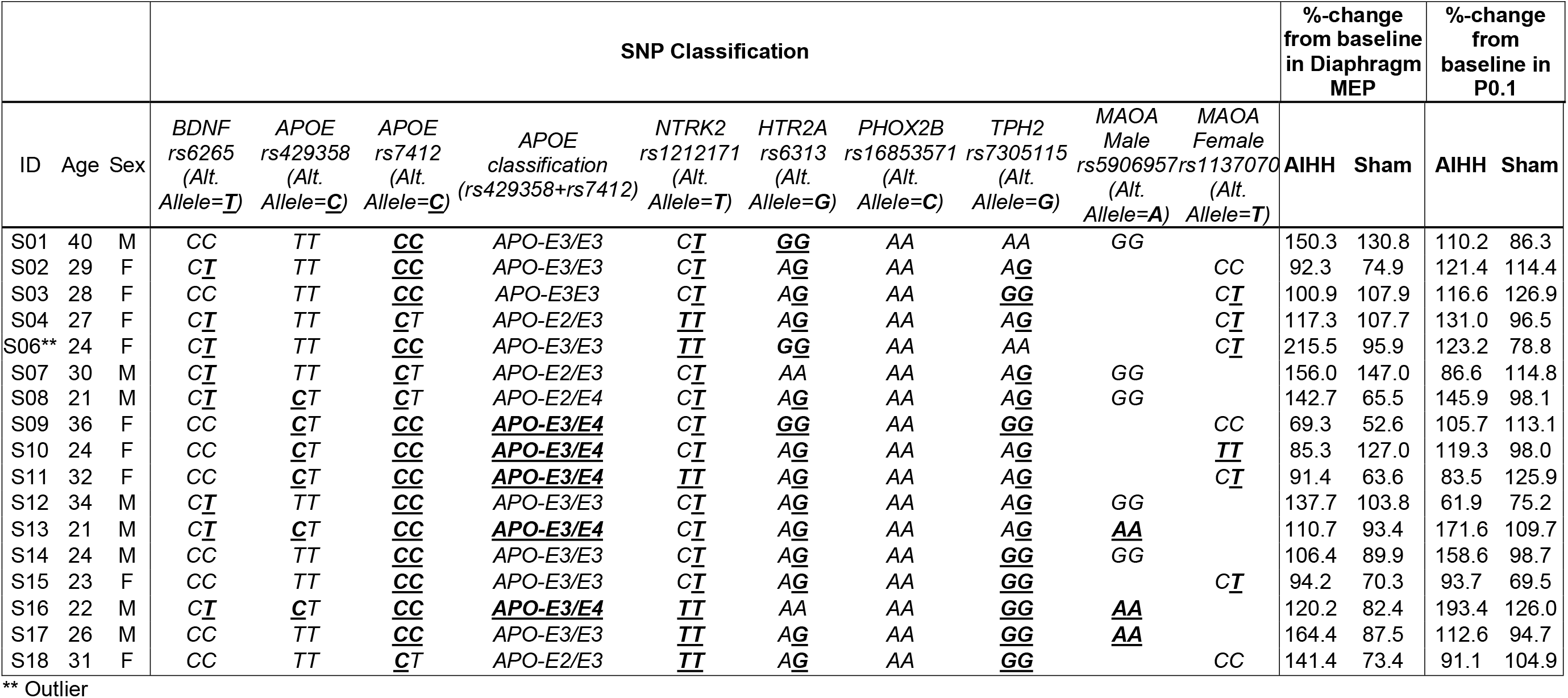
Demographics and SNP genotype classification details. Includes individual participants’ %-change from baseline in diaphragm motor-evoked potential amplitudes (MEP) and mouth occlusion pressure (P_0.1_) following AIHH and Sham exposures. Genotype letters in bold and underlined text indicate dysfunctional allele.

SNPs in sex chromosomes include male monoamine oxidase A (*MAOA*, SNP ID: rs5906957 [A>G], prevalence ~36% in male [39]) and female *MAOA* gene (SNP ID: rs1137070 [C>T], prevalence ~31% in female [40]).

### DNA Extraction and Genotyping

#### Saliva collection and storage

Participants drool saliva was collected in a DNA/RNA shield-saliva collection kit (Genesee Inc.). Genomic (g) DNA from the saliva was extracted using a spin column-based DNA isolation kit (Zymo Quick-DNA Miniprep Kit Cat# D4069). Extracted gDNA was quantified via spectrophotometry (NanoDrop Model 2000C, Thermo Fisher Sci.) and sample purity was estimated by absorbance ratio of A260/A280 (sample range: ?1.8-2.0). Extracted DNA was diluted to 1ng/ul concentration and used as templates in real time quantitative polymerase chain reaction (PCRs; QuantStudio3; Applied Biosystems). A 5’ to 3’ exonuclease assay in TaqMan (Applied Biosystems) was used to amplify the gene SNP of interest. SNP genotyping calls were performed with TaqMan Genotyper Software (Thermo Fisher Sci. Inc). Human DNA samples with known genotype from Coriell Institute’s Medical Research Repository were used as control identifier for TaqMan Genotyper Software.

#### Genotype coding used for regression analysis

Prior to applying linear model regression for SNP loci analysis, genotypes were recoded: 1) for *BDNF*, the “T” allele number was counted; 2) for *APOE*, the number of allele “C” in 2 loci, i.e. rs429358, and rs7412 were counted, and if the number was ?3, the new variable was set to 1 (otherwise 0); 3) for *NTRK_2_*, the number of allele ‘T’; 4) for *HTR_2A_*, the number of allele ‘G’; 5) for *TPH_2_*, the number of allele ‘G’ was counted. Since *MAOA* SNP loci (male, rs5906957 and female, rs1137070) have different localizations on the X chromosome, we stratified results based on sex and analyzed them separately. Data from *PHOX_2B_* SNP (rs16853571) was omitted in the analysis due to lack of gene variation in our study sample. For SNP locus analysis, variables age and sex were considered as covariates.

### Humanized *ApoE* Knock-in Rat Experiments

Based on the observed association between *APOE_3/4_* and impaired AIHH-induced diaphragm plasticity in humans, we performed follow up experiments in adult male Sprague-Dawley rats (345-385g; Envigo, IN, USA) with homozygous knock-in humanized *ApoE3 (hApoE_3_;* ID #395, n=4) or *ApoE4* (hApoE_4_; ID #359, n=3). Neurophysiology experiments were performed in urethane anesthetized, paralyzed and ventilated rats at times consistent with human AIHH treatments (*i.e.*, active phase; 12 a.m. in rats [41]). The primary outcome measure was the amplitude of integrated phrenic nerve bursts (1-min averages), taken before, during, and 30, 60 and 90 min after exposure to an AIHH protocol comparable to that delivered to humans (15, 1 min episodes of hypercapnic-hypoxia; 1.5 min intervals). Experimental details of these neurophysiology experiments are provided in the supplemental section and elsewhere [42–44].

### Statistics

The quality of SNP genotype data was analyzed for deviations from Hardy Weinberg equilibrium using both the Exact Test and Chi-Squared Test. A single-locus analysis was used to assess the association of each SNP with treatment outcome [45]. After adjusting for age and sex, the association between %-change from baseline and SNPs was explored using a linear regression model in R software [46]. A detailed description of SNP genotype coding used for liner regression analysis is provided in the supplementary section. The association of age and sex with primary dependent variables (diaphragm MEPs and P_0.1_) were analyzed using a liner regression model.

Peak phrenic nerve burst amplitude was averaged over 1 min immediately before blood samples were taken at baseline and at 30, 60 and 90 min post-AIHH. Phrenic nerve burst amplitude was analyzed using absolute values and normalized as a percent change from baseline. Phrenic responses were analyzed using a two-way repeated measures ANOVA with Tukey’s *post-hoc* analysis (SigmaPlot, v12.0; Systat Software, San Jose, CA). Differences were considered significant when p<0.05. Data are expressed as mean ± SD.

## RESULTS

Demographics, genotype and pre to post %-change in primary dependent variables (MEP and P_0.1_) following AIHH and Sham for each participant are presented in Table 1. A detailed report of the cardiorespiratory responses during AIHH exposure in the same set of individuals is presented in a companion paper [3]. Only genetics, age, and sex effects on diaphragm MEP amplitudes and P_0.1_ are presented here; age and sex effects are presented in supplementary material.

### Gene SNPs Associated with Dysfunctional AIHH-Induced Plasticity

No departure from Hardy-Weinberg equilibria was observed within the screened autosome or sex chromosome loci. For brevity, and due to their associations with AIHH-induced plasticity, we report results in this manuscript for *BDNFval/met, APOE_4_* and *TPH_2_* SNPs. A complete summary of all SNPs and multiple regression analyses for %-change in diaphragm MEP amplitudes and P_0.1_ are provided in Tables 2A and 2B. One participant (participant ID: S06; Table 1) with *TPH_2_* homozygous major “A” allele was identified statistically (Cook’s D >4) as the most influential data point in the regression for %-change in diaphragm MEP amplitudes (Figure 2). Therefore, data from S06 was not included in any analysis except for *TPH_2_* group analysis.

**Figure 2:**
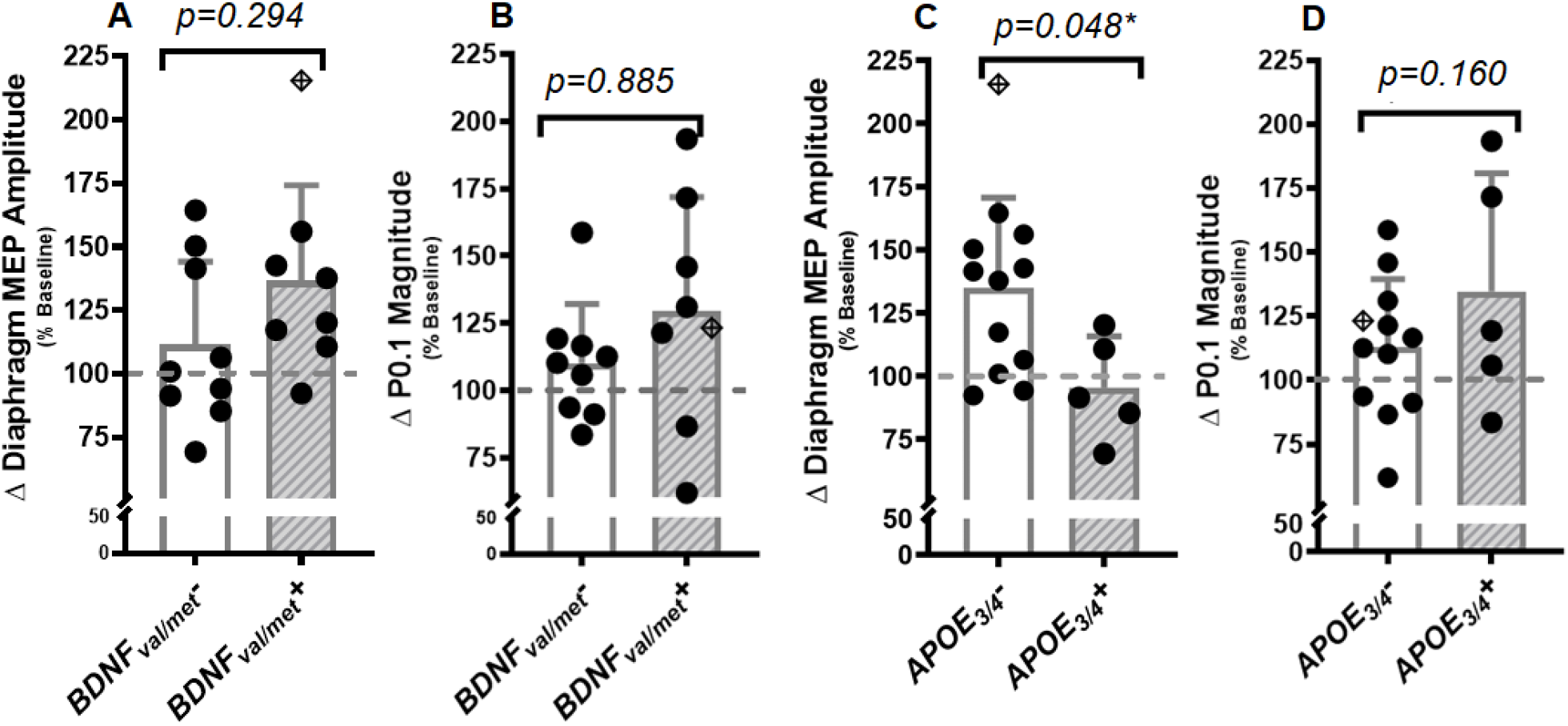
Relative (%-change from baseline) changes in diaphragm motor-evoked potential (MEP) amplitudes and mouth occlusion pressure (P_0.1_) in individuals with *BDNFval/met* (panels A and B) and *APOE_3/4_* (panels C and D) SNP. No associations were observed between individuals with *BDNFval/met* and the change in MEP amplitudes (panel A) or P_0.1_ (panel B). Individuals with dysfunctional *APOE_3/4_* allele were associated with a significantly lower AIHH-induced change in MEP amplitude (t=-2.28, p=0.048, panel C). However, no association between *APOE_3/4_* and AIHH-induced P_0.1_ responses were observed (panel D). Δ=change. *p<0.05. Results expressed as mean ± SD. 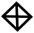 *= participant (S6) was identified as the most influential point (Cook’s D >4) in the %-change in diaphragm MEP amplitudes, therefore, the data was not included in group analyses.*

**Table 2.**
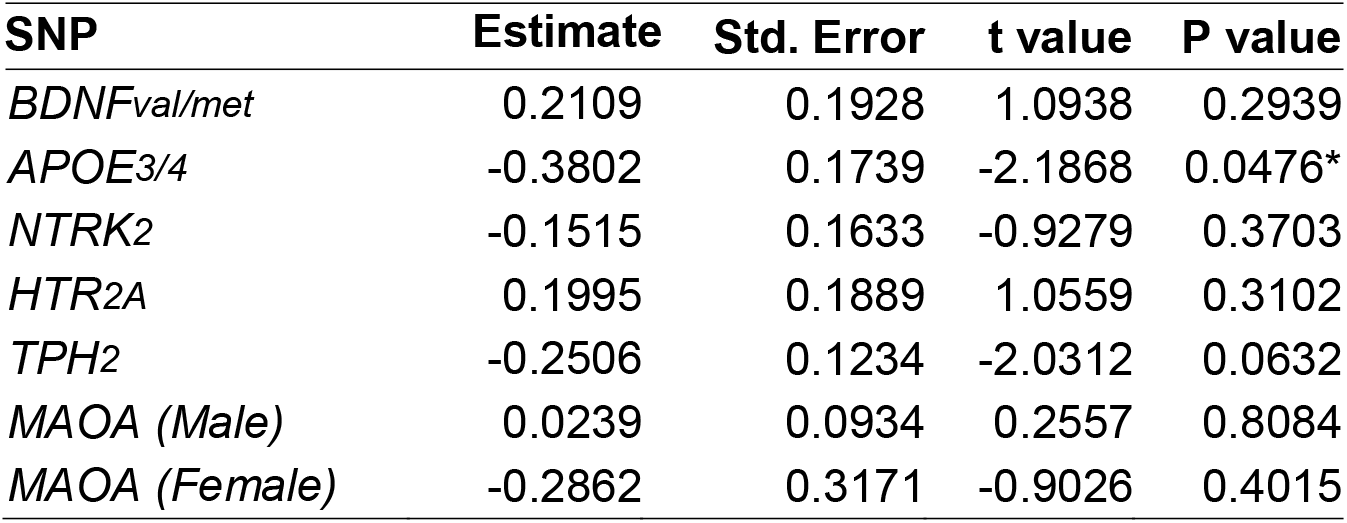
Association of SNPs with %-change from baseline in diaphragm motor-evoked potential amplitudes (MEP). *p<0.05.

#### BDNFval/met (rs6265)

Eight participants were heterozygous, and none were homozygous for the *BDNFval/met* allele. No significant difference was observed between *BDNFval/met* heterozygotes and individuals without *BDNFval/met* for %-change in diaphragm MEP amplitudes (Figure 2A; Table 2, p=0.290, t=1.090) or P_0.1_ (Figure 2B; Table 3, p=0.885, t=0.150).

**Table 3.**
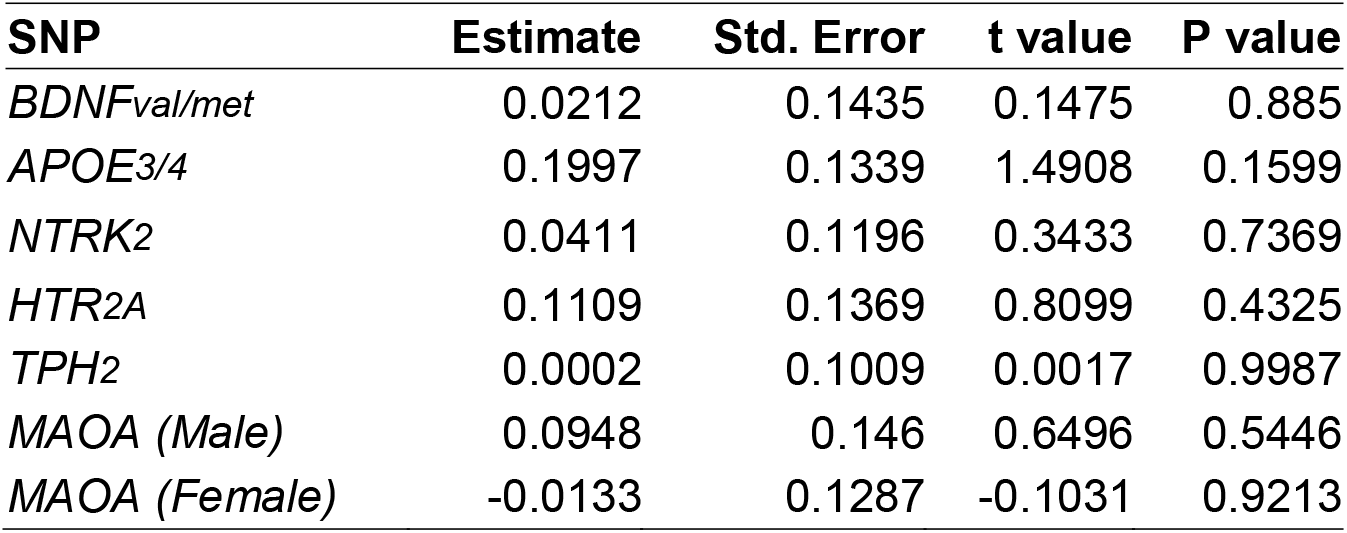
Association of SNPs with %-change from baseline in mouth occlusion pressure in 0.1 seconds (P_0.1_). *p<0.05.

| SNP | Estimate | Std. Error | t value | P value |
| --- | --- | --- | --- | --- |
| *BDNF<sub>val/met</sub>* | 0.0212 | 0.1435 | 0.1475 | 0.885 |
| *APOE<sub>3/4</sub>* | 0.1997 | 0.1339 | 1.4908 | 0.1599 |
| *NTRK2* | 0.0411 | 0.1196 | 0.3433 | 0.7369 |
| *HTR2A* | 0.1109 | 0.1369 | 0.8099 | 0.4325 |
| *TPH2* | 0.0002 | 0.1009 | 0.0017 | 0.9987 |
| *MAOA (Male)* | 0.0948 | 0.146 | 0.6496 | 0.5446 |
| *MAOA (Female)* | -0.0133 | 0.1287 | -0.1031 | 0.9213 |

#### APOE (rs429358 and rs7412)

Five participants were heterozygous for *APOE_4_* (i.e., *APOE_3/4_);* none were homozygous for *APOE_4_*. The *APOE_3/4_* genotype was associated with diminished %-change in diaphragm MEP amplitudes following AIHH (Figure 2C, Table 2, p=0.048, t=-2.187). The %-change in diaphragm MEP amplitudes was 38% lower in individuals with *APOE_3/4_ (APOE_3/4_+) versus* individuals carrying other allelic *APOE* isoforms (e.g., *APOE_3/4_*). In contrast, no significant association between *APOE_3/4_+* and %-change in P_0.1_ was observed (Figure 2D; Table 3, p=0.159, t=1.490).

#### TPH_2_ (rs7305115)

Two participants were homozygous for the *TPH_2_* major “A” allele (participant ID: S01 and S06), 8 participants were heterozygous and 7 homozygous for the dysfunctional minor “G” allele. Although not statistically significant, there was a marginal association between the presence of at least 1 “G” allele and %-change in diaphragm MEP amplitudes (p=0.063, t=-2.030). The coefficient of the *TPH_2_* gene was −0.251, meaning responses were 25.1% lower than average with 1 “G” allele. This effect was primarily influenced by the outlier participant (S06) who was homozygous for “A” allele. No association was observed between *TPH_2_* locus variants and P_0.1_ (p=0.990, t=0.002).

### Age-Sex Dimorphism in Diaphragm MEPs

No significant relationship was found between age and %-change in diaphragm MEP amplitude following AIHH (Figure 4A; r=0.08, 95% CI= −2.47 to 3.32, p=0.758). No significant differences in diaphragm MEP amplitude change were observed with age in males (Figure 4B; r=0.24, 95% CI= −1.18 to −0.4.24, p=0.217) or females (Figure 4B; r=-0.01, 95% CI= −5.75 to 4.38, p=0.752). However, males had significantly higher %-change in diaphragm MEP amplitudes *versus* females, regardless of age (mean difference=37±10.8%, F=12.17, p=0.004).

### Age-Sex Dimorphism in P_0.1_

A negative correlation was observed between %-change in P_0.1_ and participant’s age, despite the limited age range included in this study (Figure 4C; r=-0.64, 95% CI=-0.85 to-0.23, p=0.007). Each year of increasing age corresponded to a 3.9% decrease in P_0.1_ response. The decline in P_0.1_ with age was explained by male (Figure 4D; r=-0.73, 95% CI= −0.95 to −0.07, p=0.036) *versus* female responses (Figure 4C; r=-0.29, 95% CI= −0.83 to −0.52, p=0.480) to AIHH. Regression slope (F=1.77, p=0.210) and intercept (F=1.5, p=0.240) for %-change in P_0.1_ were not significantly different between males and females.

### Humanized *ApoE* Knock-In Rats and AIHH Induced Phrenic Long-Term Facilitation

Figure 3A shows average phrenic nerve burst amplitudes during and following AIHH. Baseline phrenic nerve amplitudes were not different between groups (*hApoE_3_:* 0.023±0.007 V; *hApoE_4_:* 0.022±0.013 V). On the other hand, AIHH elicited significant phrenic long-term facilitation in *hApoE_3_* (p=0.025 vs. baseline), but not in *hApoE_4_* rats (p=0.995). A significant interaction between genotype and time post-AIHH was observed in phrenic long-term facilitation magnitude (Figure 3B; F=5.93, p=0.007). AIHH-induced phrenic long-term facilitation in *hApoE_3_* rats was significantly greater than *hApoE_4_* at 30 min (p=0.004), 60 min (p=0.002) and 90 min (p<0.001) post-AIHH. Arterial CO2 partial pressures at baseline (*hApoE_3_:* 43.9 ± 1.5 mmHg; *hApoE_4_:* 45.7±1.2 mmHg) and 90 min post-AIHH (*hApoE_3_:* 44.4±1.6 mmHg; *hApoE_4_:* 46.2±0.4 mmHg) were not different.

**Figure 3:**
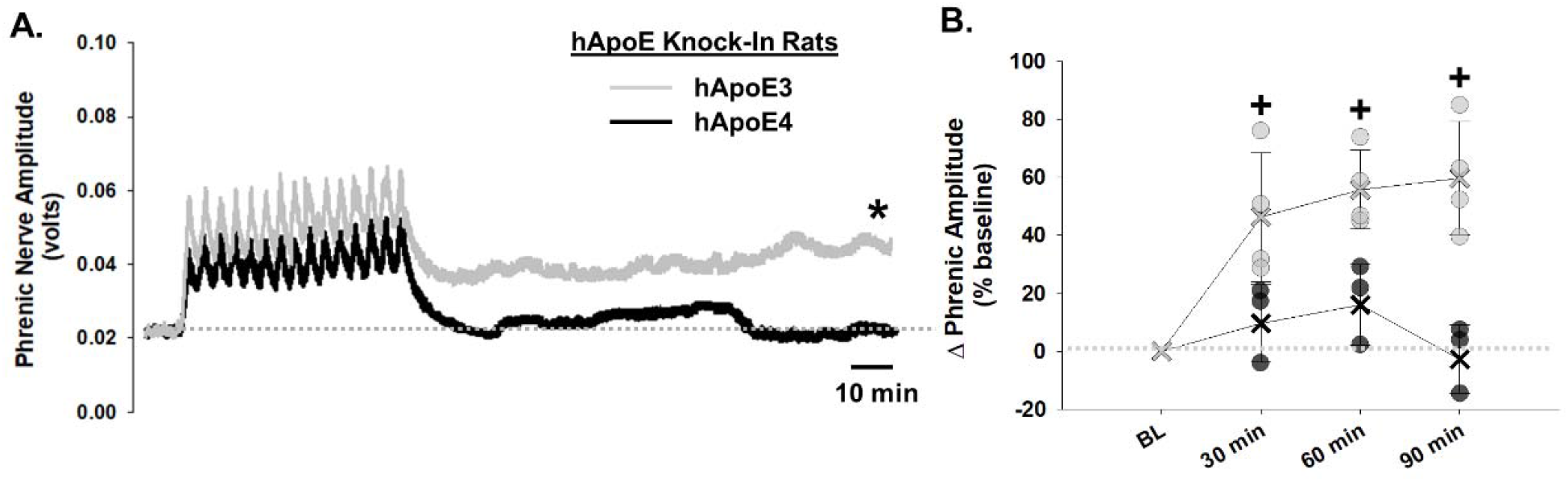
AIHH elicits phrenic long-term facilitation in hApoE_3_ but not hApoE_4_ knock-in rats. Panel A shows average traces of phrenic nerve amplitude for hApoE_3_ (n=4; gray) and hApoE_4_ (n=3; black) knock-in rats, *p<0.050 vs baseline. Panel B phrenic burst amplitude (%- change from baseline) in hApoE_3_ (gray circles) and hApoE_4_ (black circles) rats, +p<0.005 *versus* hApoE_4_. Δ=change. Results expressed as mean ± SD.

**Figure 4:**
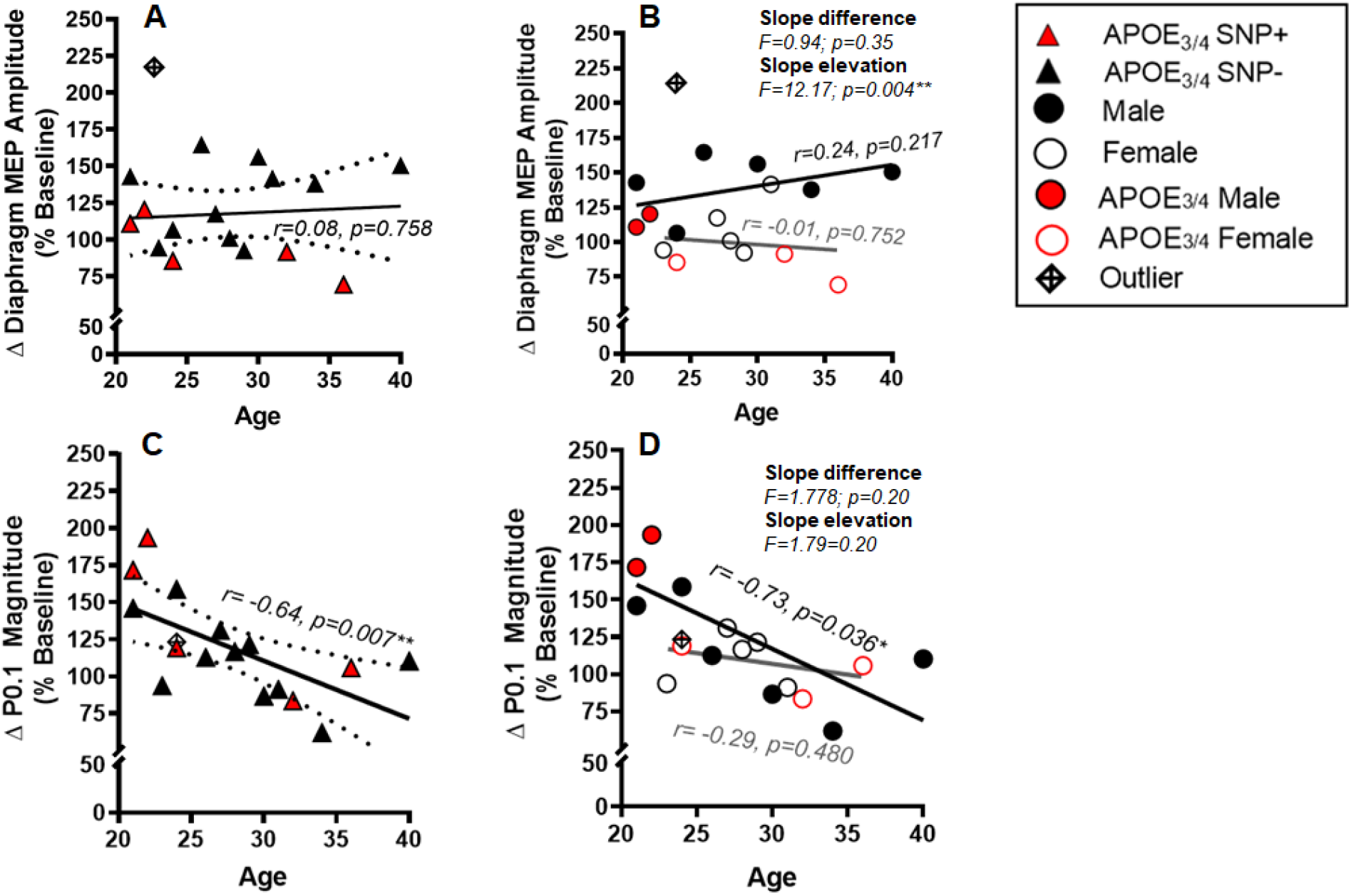
Relationship between age and sex on the magnitude (%-change from baseline) of change in diaphragm motor-evoked potential amplitudes (MEP, panels A and B), and mouth occlusion pressure in 0.1 seconds (P_0.1_, panels C and D) following AIHH. No association between age and the magnitude of change in diaphragm MEP amplitudes was observed (panel A). Regardless of age, males (black line, panel B) had significantly greater responses in MEP amplitudes *versus* females (gray line, panel B). The magnitude of change in P_0.1_ reduced significantly with age (panel C); however, the decline was more pronounced in males (r=-0.73, p=0.036, black line, panel D) *versus* females (r=-0.29, p=0.480, gray line, panel D). Δ=change. *p<0.05. Results expressed as mean ± SD. 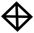 *= participant (S6) was identified as the most influential point (Cook’s D >4) in the %-change in diaphragm MEP amplitudes, therefore, the data was not included in group analyses.*

## DISCUSSION

We investigated the role of genetics, age and sex on AIHH-induced respiratory motor plasticity of both cortical (presumably volitional) diaphragm MEPs and brainstem automatic (P0.1) neural pathways in healthy adults. We report increased diaphragm MEP amplitudes following AIHH are diminished in people heterozygous for the *APOE_4_* allele and unaffected in *BDNFval/met* heterozygotes. Regardless of age, the %-change in diaphragm MEP amplitudes following AIHH is greater in males *versus* females, whereas sex does not influence the magnitude of change in P_0.1_. Finally, despite the limited age range in this study (20-40 years), there was a negative correlation between age and P_0.1_ facilitation. Neurophysiological experiments in *hApoE_3_* and *hApoE_4_* knock-in rats confirmed a causal relationship between *hApoE_4_* genotype and impaired phrenic motor plasticity.

### SNPs and AIH/AIHH Induced Plasticity

To investigate SNPs that influence AIH/AIHH-induced respiratory motor plasticity, a panel of genes was assessed chosen based on their known links to phrenic motor plasticity in rodents, including SNPs linked to serotonin synthesis (*TPH_2_*), clearance (*MAOA*), or receptors (*HTR_2A_*), a key neurotrophic factor (*BDNF*), and its high affinity receptor (*NTRK_2_*), as well as chemoreceptor function (*PHOX_2B_*). A seventh gene, *APOE_4_* was added to the panel due to its association with impaired neuroplasticity [17], including AIH-induced phrenic long-term facilitation [47].

No association was found between 6 gene SNPs and AIHH-induced respiratory motor plasticity in the humans studied here. Tryptophan hydroxylase-2 (*TPH_2_*) is the rate limiting enzyme for serotonin synthesis [36]; presence of a “G” allele in exon 7 of the *TPH_2_* gene is associated with reduced serotonin bioavailability [35, 48]. An apparent (but. not significant; p=0.063) ~25% diminished response in the presence of 1 *TPH_2_* “G” allele requires further study.

Since BDNF is both necessary and sufficient for AIH-induced phrenic motor plasticity in rats [14], we hypothesized that the dysfunctional *BDNFval/met* allele undermines plasticity. *BDNFval/met* is a common missense single nucleotide C>T polymorphic mutation at codon 66 of *BDNF* gene, resulting in amino acid methionine (Met) substituting valine (Val). *BDNFval66met* or *BDNFval/met* mutation, impairs the pro-domain region of BDNF protein, disrupting the normal trafficking of mature BDNF from neuron soma to dendrites [49–51]. This dysfunctional *BDNF* SNP is associated with reduced exercise-induced plasticity and functional recovery in people with spinal cord injury or traumatic brain injury [31, 52, 53]. However, contrary to our hypothesis, no association between *BDNFval/met* mutation and AIHH-induced respiratory motor plasticity was found (Figure 2A). We speculate that in healthy adults, one fully functional allele is sufficient to meet physiological demands and/or enable adequate responses to certain physiological stimuli, such as AIHH. Since no participants had homozygous *BDNFval/met* mutation, we cannot rule out an association between homozygous *BDNFval/met* and respiratory motor plasticity.

*APOE* is a triglyceride rich low-density lipoprotein that facilitates lipid transport between cells. *APOE* is highly expressed in the central nervous system, with 3 common human isoforms (E2, E3 and E4) [54]. With respect to neuroplasticity, the T to C nucleotide substitutions at *APOE* loci (*APOE_4_*) leads to arginine substitutions in the 112 and 159 positions (SNPs rs429358 and rs7412), and is the most consequential SNP mutation for neuroplasticity. Homozygous *APOE_4_* allele is present in 11-14% of people, whereas heterozygous *APOE_3/4_* allele is found in about 15-25% of people [27, 28]; In this group of study subjects, we observed a slightly higher percentage of *APOE_3/4_* heterozygotes (~29%), which may be attributed to our small sample size. Individuals with the *APOE_4_* allele experience diminished motor recovery following spinal cord injury *versus* other *APOE* alleles [25]. *APOE_4_* protein isoform has been hypothesized to impair AIH-induced plasticity [47] as it reduces NMDA and AMPA receptor recycling in the post-synaptic membrane, and limits BDNF availability. A recent study in transgenic mice with knock-in *hApoE_4_* suggested that APOE_4_ protein isoform is associated with impaired AIH-induced respiratory motor plasticity [47], consistent with our observation that at least 1 dysfunctional *APOE_4_* allele was associated with 38% reduction in AIHH-induced diaphragm MEP facilitation. Thus, stratifying participants based on Mendelian randomization of known genetic risk factors may be critical for success of large phase II and III clinical trials investigating the efficacy of AIH/AIHH [56].

### Causal link between *APOE_4_* on AIHH-induced respiratory motor plasticity

To demonstrate a causal link between *APOE_4_* and AIHH-induced respiratory motor plasticity, we performed neurophysiology experiments in *hApoE_4_* and *hApoE_3_* knock-in rats using a nearly identical AIHH protocol to humans (15, 1-minute episodes of hypercapnic-hypoxia during the night, or the active phase for rats). Whereas rats with *hApoE_3_* manifested robust AIHH-induced phrenic long-term facilitation (~60% increase at 90 min post AIHH), *hApoE_4_* rats failed to express significant plasticity. Thus, *APOE_4_* undermines AIHH-induced respiratory motor plasticity in rats. Our data support an earlier report by Strattan and colleagues [47] where *hApoE_4_* mice failed to express AIH-induced respiratory plasticity, despite study differences such as species (mice *versus* rats), plasticity-inducing protocol (AIH *versus* AIHH) and time of day (rest *vs* active phase).

Although the mechanistic link between a dysfunctional *APOE_4_* allele and reduced spinal plasticity is not yet known, we suggest a few plausible hypotheses. APOE_4_ protein isoform converts microglia to a pro-inflammatory phenotype [26], which may undermine phrenic motor plasticity [55]. Further, the observation that *hApoE_4_* mice exhibit more extensive perineuronal nets after spinal cord injury [47] suggests an alternate mechanism, and suggests a distinct therapeutic target to mitigate the dysfunctional effects of *APOE_4_* genotype. Future studies investigating *APOE_4_* induced pathophysiology may reveal additional targets to unlock AIHH induced neuroplasticity in *APOE_4_* carriers.

Unlike the association of *APOE_3/4_* and *TPH_2_* SNPs with reduced diaphragm MEP responses following AIHH, no similar association was found between these genotypes and P_0.1_. This difference could be due to distinctions in the neuronal pathways utilized with transcranial magnetic stimulation (reflecting volitional control of breathing) *versus* automatic (bulbospinal) pathways to phrenic motor neurons and/or the correlation between participant’ age and P_0.1_ facilitation (see below), which likely obscured the influence of genetic factors.

### Age-Sex Dimorphism in AIHH Induced Plasticity

Decades of rodent work demonstrate a link between age, sex and AIH-induced phrenic motor plasticity [18, 19, 56, 57]. Although our results are generally consistent with prior observations in rats, there were some interesting differences.

#### Diaphragm MEP responses

We observed that in healthy adults, regardless of age, corticospinal plasticity (i.e., diaphragm MEPs) was significantly greater in males versus females (mean difference=37±10.8%). Sex differences in the neural control of breathing have been observed during ventilatory challenges [58, 59] and the capacity for respiratory neuroplasticity [60, 61]. These sex differences could be caused by ovarian hormones that affect neurotransmission. In rats, hippocampal long-term potentiation is induced more readily in males *versus* females due to excitatory effects of testosterone [62, 63]. In females with normal menstruation, circulating progesterone reduces cortical excitability [64, 65]. During the luteal phase of menstrual cycle (high progesterone), increased inhibition and decreased facilitation of TMS responses are observed, which is indicative of increased GABAergic effects from progesterone metabolites [65]. In contrast, there is increased cortical facilitatory activity during the mid-follicular phase of the menstrual cycle (low progesterone, high estrogen). Thus, our results are in line with previous literature.

#### P0.1 responses

A significant decrease in AIHH-induced P0.1 plasticity was observed with increasing age; each year of age in the range studied (20-40 years) led to a fall in P0.1 plasticity of ~3.9%. This age-related drop was more pronounced in males than females. Negative pressure generation in 0.1 seconds of an occluded inspiration reflects respiratory neuromechanical drive prior to influences from breath-related sensory feedback, such as from lung or chest wall receptors [22]. Explanations for diminished AIH/AIHH-induced neuroplasticity with age observed in the present study include: 1) decreasing sex hormone (testosterone/estrogen) levels [19, 66]; 2) diminished serotonergic function [18] and/or 2) increased extracellular CNS adenosine levels [67, 68].

Since changes in P0.1 reflect automatic control of breathing, it may be more equivalent to rodent phrenic long-term facilitation versus MEPs. In rats, phrenic long-term facilitation decreases as males reach middle-age [19], but increases in middle-aged females (when normalized for stage of the estrus cycle) [69]. Estrogen suppresses pro-inflammatory microglial activities [70] and even mild inflammation impairs phrenic long-term facilitation [55, 71, 72]. Testosterone is necessary for phrenic long-term facilitation in males because it is a substrate for aromatase-dependent CNS estrogen formation [66]. In male rats, testosterone peaks at ~2-6 months of age, equivalent to ~18-40 years in humans [73–75], which is then followed by a gradual decline, similar to human males in the ~40-60 year age range [76, 77]. Since the age of our participants ranged from 20-40 years, reduced serum sex hormone levels are unlikely to explain variance in P0.1 responses; furthermore, the %-change in P0.1 was not significantly different between sexes in this study. Adenosine is another major regulator of AIH-induced phrenic motor plasticity in rats [78, 79]. Extracellular adenosine levels in the central nervous system increase with age – greater adenosine-dependent inhibition of phrenic motor plasticity may occur [80], potentially explaining reduced P0.1 plasticity with age in our study.

## LIMITATIONS

Rather than more common “omics” approaches, we selected a panel of 7 genes and 9 SNPs to investigate as a potential biomarker based on known roles of the relevant molecules in AIH-induced respiratory motor plasticity, and the relative penetrance of the SNPs in humans. Although this list does not include all SNPs that could affect AIH/AIHH-induced plasticity, we verify that genetic factors regulate AIHH-induced plasticity in humans, particularly *APOE_4_.* This is the first study to link *APOE_4_* with spinal, respiratory motor plasticity in humans. Due to the number of potential SNPs that could be investigated, adequate correction for multiple comparisons will remain a challenge. Further, it is important to increase the age range studied beyond 40 years and to extend investigations to people living with disease or injury.

## CONCLUSIONS

We provide evidence that the *APOE_4_* allele, age and sex are important biological determinants of AIHH-induced respiratory motor plasticity in humans. The presence of one dysfunctional *APOE_4_* allele undermines cortico-spinal respiratory motor plasticity. Experiments using humanized *APOE_4_* knock-in rats support a causal relationship between *APOE_4_* and impaired AIHH-induced respiratory motor plasticity. Contrary to our original hypothesis, no evidence was found for diminished plasticity in individuals with *BDNFval/met* mutations, although no homozygous subjects were included in this analysis. Regardless of age, males exhibited greater AIHH-induced cortico-spinal plasticity *versus* females; conversely, AIHH-induced plasticity in P0.1 is negatively associated with increasing age – an effect that is more pronounced in males than females. Thus, age, sex and genetic factors should all be considered when attempting to differentiate responders from non-responders in clinical trials investigating therapeutic use of AIH/AIHH in individuals with spinal cord injury or other neurological conditions. With such information in hand, it may be possible to refine rehabilitation protocols and/or provide individualized treatment strategies.

## Supporting information

Nair et al. Supplemental Data

## ACKNOWLEDGEMENTS

The authors thank Carter Lurk and Patrick Argento for their technical assistance with sample management and genotyping; both have reviewed the final manuscript and provided permission for this acknowledgement.

