## Supplementary material for "APOE4, Age & Sex Regulate Respiratory Plasticity Elicited By Acute Intermittent Hypercapnic-Hypoxia": Nair et al. Supplemental Data

Department of Physical Therapy

College of Public Health and Health Professions

1225 Center Drive

Gainesville, FL, USA 32611

Funding	3

ADDITIONAL PROTOCOL AND METHODS	4

AIHH Exposure Protocol	4

Respiratory Neuroplasticity Outcomes	5

Diaphragm Motor-Evoked Potentials (MEP)	5

Electromyography	5

Mouth Occlusion Pressure at 0.1s *(P0.1)*	6

Genotype Coding used for Regression Analysis	6

Rodent Neurophysiological Recordings	6

SUPPLEMENTAL RESULTS	7

Rodent Neurophysiological Experiments	7

Effects of Age and Sex on Ventilation	7

ADDITIONAL DISCUSSION	8

Age-Sex Dimorphism in Ventilation Following AIHH	8

SUPPLEMENTAL REFERENCES	9

SUPPLEMENTAL FIGURE & TABLE LEGENDS	10

Figure E1	12

Figure E2	13

Table E1	15

Table E2	16

**Funding:** National Institutes of Health (HL148030 & HL149800), Craig H. Neilsen Foundation (JW), Department of Defense SCIRP Clinical Trial Award and UF Brain and Spinal Cord Injury Trust Fund (BSCIRTF).

**ADDITIONAL PROTOCOL AND METHODS**

**AIHH Exposure Protocol**

Participants were exposed to either AIHH or Sham in a random order on 2 days (separated by ≥ 3 days of washout period). Participants received exposures in a comfortable seated position. Each session lasted ~2 hours and consisted of 5-10 min of baseline eupneic breathing followed by ~45 min of experimental gas exposures. These exposures consisted of AIHH (15, 1-min hypercapnic-hypoxic episodes with 1.5 min intervals breathing room air) and normocapnic-normoxia (Sham control). Total exposure duration was 15 min (Figure E1); total protocol duration was 36 min. At the end of the last exposure, the participant was allowed a 30 minutes of comfort break. Eupneic breathing was continuously measured between 30-60 min post-exposure. To ensure participant safety, level of discomfort and cardiorespiratory responses were monitored throughout the session. During baseline and post-exposure eupneic breathing measurements, as well as during AIHH (or Sham) exposures, participants wore a face mask (7450 V2, Hans Rudolph Inc.) attached to a two-way non-rebreathing valve (2700, Hans Rudolph Inc.). The inspired limb of the valve was connected to a calibrated pneumotachograph (3818, Hans Rudolph Inc.) and a 3-way stop-cock (2100, Hans Rudolph Inc.); these items were hidden from participant’s sight. One arm of the stop-cock received room air; the other received gas from a large Douglas bag (1196, VacuMed). The Douglas bag was filled with ~9.5% O2 and 4.5% CO2 (balance N2) to deliver AIHH. Participants breathed ambient air during Sham exposures. Inspired and expired gases were sampled at the mouth using a calibrated O2/CO2 gas analyzer (GEMINI, CWE Inc.; Ardmore, PA, USA). Mouth pressure was obtained from a port in the facemask connected to a differential pressure transducer (113253, Hans Rudolph Inc.). Inspired flow was corrected for inspired gas composition effects on airflow resistance [48] using a continuously adapting correction factor [49]. Tidal volume (VT), breathing frequency (fb) and minute ventilation ($\dot{\text{V}}$I) were derived from inspired flow and volume.

**Respiratory Neuroplasticity Outcomes**

***Diaphragm motor-evoked potentials (MEP).*** Transcranial magnetic stimulation was performed over the left motor cortex with the aim of stimulating areas involved in activation of the right diaphragm, as described by Maskill et al., 1991 [1]. The vertex of the skull was identified by intersection between nasion to inion (sagittal plane) and tragus to tragus (coronal plane). Stimuli were delivered using a handheld 70-mm figure-of-eight coil (P/N 3190-00) powered by a magnetic stimulator (200-2, Magstim, Whitland, UK). The optimal coil position was identified for each individual by a cortical mapping procedure [2], and the site was marked directly on the scalp or over a cloth treatment cap (MagVenture; Farum, Denmark). Respiratory movements were monitored using piezoelectric transducer device (PneumoTrace II, UFI; Morro Bay, CA, USA) securely placed around the abdomen at the level of the umbilicus. The magnetic stimulus was delivered at end-expiration when the participant was at passive functional residual capacity. Resting motor threshold was identified as the lowest stimulation intensity that elicited a MEP peak-to-peak amplitude ≥ 50 μV in 3 consecutive stimulations. All subsequent stimuli were performed at an intensity equivalent to 120% of the resting motor threshold. Evoked potential amplitudes were assessed at baseline and 60 min post-exposure to AIHH, AIH and Sham.

***Electromyography.*** Magnetic stimulation evoked motor potentials were recorded from the right costal diaphragm using self-adhesive Ag/AgCl electrodes (6801, The Prometheus Group; Dover, NH, USA) placed on the chest wall in bipolar arrangement (~3 cm apart) between the sixth and eighth intercostal spaces along the anterior-axillary line. The ground electrode was placed on the acromion process of the scapula. Skin was lightly abraded and cleaned before electrode placement. Surface recordings of diaphragm EMG in response to TMS has been previously validated [3] [4] and reliability established [2]. Data were digitized (PowerLab 8/35, ADInstruments) and recorded at 10 kHz for EMG using data acquisition software (LabChart V8.1, ADInstruments).

***Mouth occlusion pressure at 0.1 sec (P0.1).*** Mouth occlusion pressure at 0.1s after the onset of an occluded inspiration is an indicator of neuromechanical drive to breath prior to breath-related sensory feedback [5]. Participants sat quietly and breathed spontaneously through the breathing circuit. Once in every ~5-20 breaths, the inspired limb of the circuit was briefly occluded prior to inspiration (not visible to participant). A minimum of 5 trials were performed. Mouth occlusion pressure was assessed at baseline and between 30-60 min post-exposure to AIHH and Sham.

**Rodent Neurophysiological Recordings**

Terminal neurophysiological experiments were performed as previously described [6-8]. Rats were induced with 3.0% isoflurane in 3 L/min O2, weighed, and transferred to a heated surgical table where anesthesia was continued (3.0% isoflurane) at inspired fraction (Fi) O2 at 0.6, balance N2. Higher FiO_2_ is maintained in this terminal preparation to sustain blood O_2_ levels in physiological range (PaO_2_ >150mmHg) and prevent ventilation/perfusion mismatch [9, 10]. A tight titration of acid-base balance is maintained to ensure the arterial pH remains at ~7.4 and PaCo2 within ±1.5mmHg range. In addition, core body temperature was also kept between 37.0 ± 1.0˚C with an external heat pad to maintain constant metabolic rate. Rats were tracheotomized and a polyethylene catheter (I.D. 1.67 mm) was inserted into the trachea for mechanical ventilation (0.07 mL/10g bw; 72 breaths/min; VentElite small animal ventilator; Harvard Apparatus, Holliston, MA, USA). End-tidal PCO2 (PETCO2) was continuously monitored (Capnogard, Novametrix, Wallingford, CT) and maintained by adjusting FiCO2, as needed.

Anesthesia was converted intravenously to urethane (2.1g/kg at 6mL/h) through tail vein catheter (24 gauge, Surflo, Elkton, MD). Anesthetic depth was assessed by toe-pinch withdrawal reflex. Supplemental anesthetic was infused as required. Rats were bilaterally vagotomized at the mid-cervical level to prevent phrenic nerve entrainment with the ventilator. The right femoral artery was cannulated with polyethylene tubing (I.D. 0.58 mm) to monitor blood pressure (TA-100 Transducer Amplifier, CWE, Inc.) and sample blood gases (ABL 90 Flex, Radiometer, OH, US). Fluids were administered intravenously to maintain acid-base balance (1.5 mL/h; 1:4 of 8.4% Na2CO3 in standard lactated Ringer’s solution). Rats were paralyzed with pancronium bromide (2 mg/kg; Sigma-Aldrich, St. Louis, MO).

The left phrenic nerve was isolated near the brachial plexus, cut distally, and de-sheathed to record respiratory neural activity using custom suction electrodes. Nerve activity was amplified (10K, A-M systems, Everett, WA), filtered (band-pass 0.3-5 kHz), and digitized (CED 1401, Cambridge Electronic Design, UK). Data were rectified, smoothed (time constant, 50 ms), and analyzed using Spike2 software (Cambridge Electronic Design, v 8.20).

**SUPPLEMENTAL RESULTS**

**Rodent Neurophysiological Experiments**

Baseline conditions were established at PETCO2 2 mmHg above recruitment threshold. Arterial PCO2 was maintained isocapnic (±1.5 mmHg) with baseline blood gas values (Table E1). Baseline O2 levels (0.6 FiO2, balance N2 and CO2) were maintained for the duration of the experiments, except during the 15, 1-min hypoxic and hypercapnic episodes (Table E2; PaO2: 40-50 mmHg, 0.09 FiO2; PaCO2: 50-55 mmHg, +5-10 mmHg above baseline; 0.04 FiCO2; PETCO2: ≥ 48 mmHg). The PaO2 levels for the entire duration of the neurophysiological preparations were maintained >150mmHg [9, 10]. All rodent neurophysiology experiments were conducted at times consistent with human activity state when receiving AIHH treatments.

**Effects of Age and Sex on Ventilation**

A declining trend in tidal volume (VT) %-change was observed with increasing age (r=-0.43, 95% CI=-0.75 to 0.07, p=0.09). For each year of increase in age, VT fell by 1.34%. Males had a significant negative correlation in VT %-changes with age (r=-0.71, 95% CI= -0.9 to -0.01, p=0.05*, Figure E2A), while females showed no trend (r=-0.14, 95% CI= -0.74 to 0.56, p=0.71, Figure E2B). The %-change in fb was not related to age (r=0.24, 95% CI=-0.27 to 0.65, p=0.35), or males (r=0.21, 95% CI=-0.58 to 0.79, p=0.62, E2C) versus females (r=0.29, 95% CI=-0.46 to 0.80, p=0.44, E2D), as indicated by the slope (F=0.34, p=0.56) and intercept (F=3.01, p=0.10).

The %-change in minute ventilation ($\dot{\text{V}}$I) did was not significantly affected by age (r=-0.35, 95% CI=-0.71 to 0.16, p=0.17). $\dot{\text{V}}$I %-change decreased significantly with age in males (r=-0.73, 95% CI=-0.95 to -0.05, p=0.04*, E2E), but not females (r=0.16, 95% CI=-0.56 to 0.75, p=0.67, E2F). Each year increase in age led to a fall in $\dot{\text{V}}$I %-change of 1.21% in males versus 0.35% in females. However, male versus female differences in slope (F=3.07, p=0.10) and intercept (F=2.6, p=0.12) were not significant.

**ADDITIONAL DISCUSSION**

**Age-Sex Dimorphism on Ventilation Following AIHH**

As the age of the participants increased VT response to AIHH exposure trended to decrease by 1.34% per year of age. This apparent age-related decline in VT response was primarily driven by males, with no significant effects of age in females. Similarly, the AIHH induced change in $\dot{\text{V}}$I decreased 1.21% for each year of age in males, but there was no significant correlation in females. No significant differences in breathing frequency were observed with age or between sexes. The decline in VT and $\dot{\text{V}}$I responses with age is consistent with the decline in P0.1 response, indicative of a possible spinal adenosine mediated effect [11-13]. Interestingly, age- related decline in VT response were not significant in females.

**SUPPLEMENTAL FIGURE & TABLE LEGENDS**

**Figure E1**: **Schema of experimental design for acute intermittent hypercapnic-hypoxia (AIHH) and normoxic (Sham) exposures.** Eupneic breathing was continuously measured between 30-60 min post-exposure. During AIHH exposure the end tidal (PET) O2 was intermittently lowered to ~60mmHg and PET CO2 raised to ~45mmHg. The magnitude of corticospinal drive and neuromechanical automatic drive was evaluated at baseline and 60 min post-exposure to AIHH and Sham. Corticospinal drive was assessed using diaphragm motor-evoked potential amplitudes (MEP) and neuromechanical automatic drive was assessed using mouth occlusion pressure in 0.1 seconds (P0.1).

**Figure E2: Relationship between age and sex on the magnitude (%-change from baseline) in tidal Volume (∆VT, panel A, B), breathing frequency (∆fb, panel C, D), minute ventilation (∆**$\dot{\text{V}}$**I, panel E, F) following AIHH**. A declining trend was observed with age and ∆VT, that failed to reach significance (panel A, black line, panel B). The declining trend in ∆VT was primarily due to significant decline in male ∆VT (r=-0.71, p=0.047*) *versus* females ∆VT (r=-0.14, p=0.710, gray line, panel B). No significant association was observed with age on ∆fb (panel C) and ∆$\dot{\text{V}}$I (panel E) in both sexes. Δ=change. *p<0.05. Results expressed as mean ± SD.

*= participant (S6) was identified as the most influential point (Cook’s D >4) in the %-change in diaphragm MEP amplitudes, therefore, the data was not included in group analyses.*

**Table E1**: **Measurements of PaCO2, PaO2 and mean arterial pressure (MAP) during baseline (BL), 30, 60, and 90 min post-AIHH in hApoE3 and hApoE4 rodent terminal neurophysiology experiment**. Arterial PCO2 was maintained isocapnic (±1.5 mmHg) with baseline blood gas values. Baseline O2 levels (60% O2, balance N2 and CO2) were maintained for the duration of the experiments, except during the 15, 1-min hypoxic and hypercapnic episodes (Table E2; PaO2: 40-50 mmHg, 0.09 FiO2; PaCO2: 50-55 mmHg, +5-10 mmHg above baseline; 0.04 FiCO2; PETCO2: ≥ 48 mmHg).

**Table E2**: **Measurements of PaCO2, PaO2 and mean arterial pressure (MAP) during AIHH exposure.** Blood gas measurements were taken at the 1st, 8th, and 15th hypoxic hypercapnia (HH) episode (HH1, HH8, and HH15, respectively).

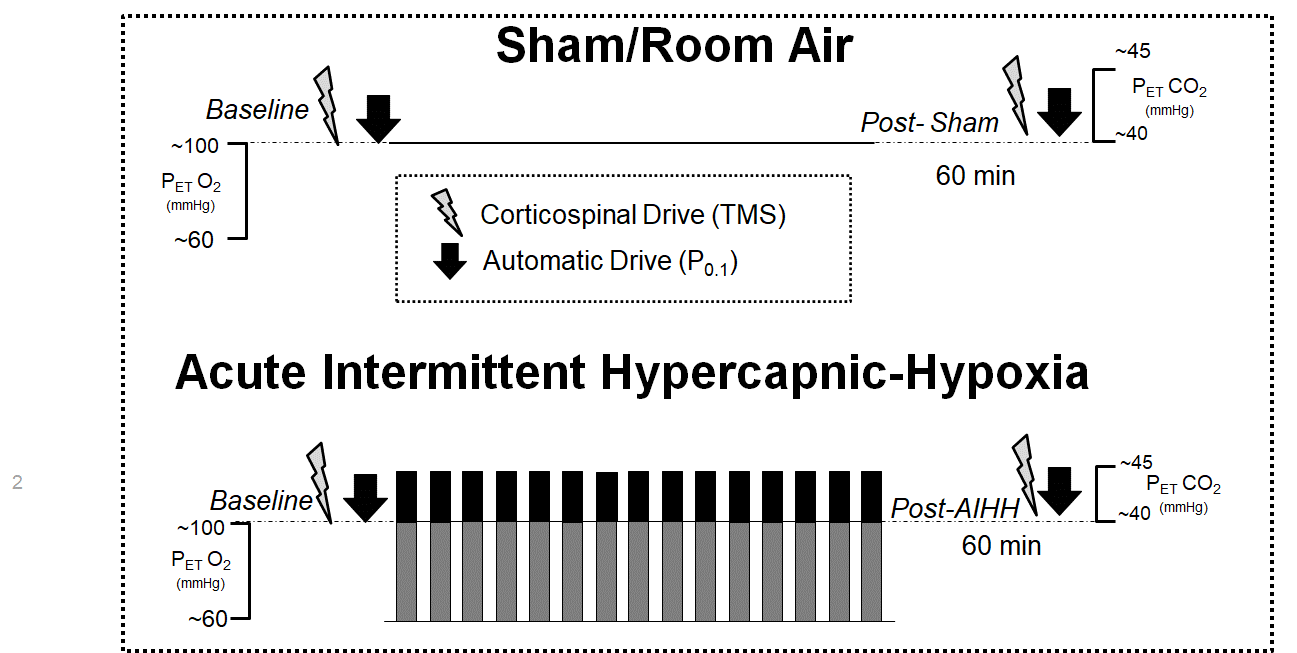

**Figure E1**: **Schema of experimental design for acute intermittent hypercapnic-hypoxia (AIHH) and normoxic (Sham) exposures.** Eupneic breathing was continuously measured between 30-60 min post-exposure. During AIHH exposure the end tidal (PET) O2 was intermittently lowered to ~60mmHg and PET CO2 raised to ~45mmHg. The magnitude of corticospinal drive and neuromechanical automatic drive was evaluated at baseline and 60 min post-exposure to AIHH and Sham. Corticospinal drive was assessed using diaphragm motor-evoked potential amplitudes (MEP) and neuromechanical automatic drive was assessed using mouth occlusion pressure in 0.1 seconds (P0.1).

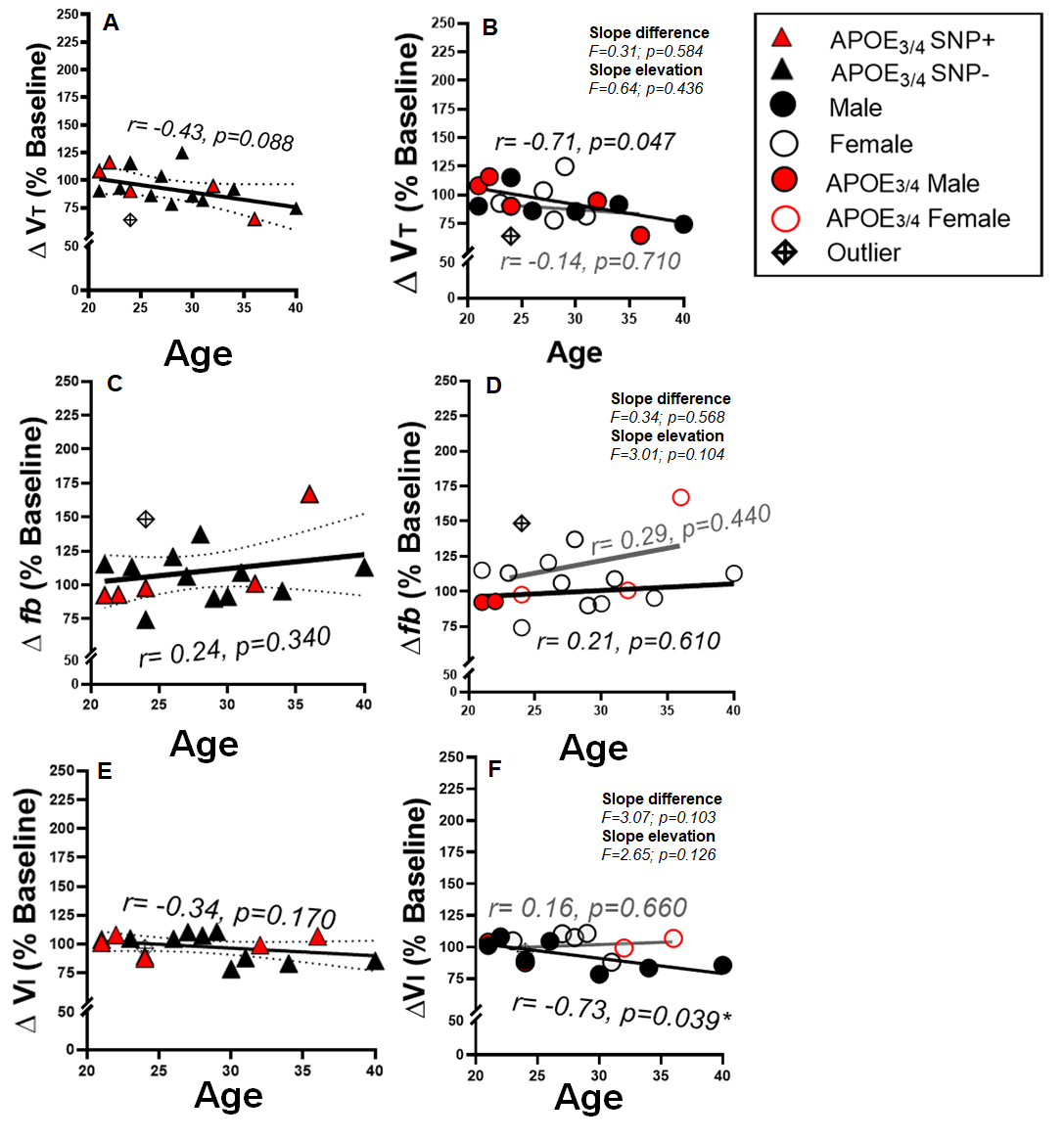
**Figure E2**: **Relationship between age and sex on the magnitude (%-change from baseline) in tidal Volume (∆VT, panel A, B), breathing frequency (∆fb, panel C, D), minute ventilation (∆**$\dot{\text{V}}$**I, panel E, F) following AIHH**. A declining trend was observed with age and ∆VT, that failed to reach significance (panel A, black line, panel B). The declining trend in ∆VT was primarily due to significant decline in male ∆VT (r=-0.71, p=0.047*) *versus* females ∆VT (r=-0.14, p=0.710, gray line, panel B). No significant association was observed with age on ∆fb (panel C) and ∆$\dot{\text{V}}$I (panel E) in both sexes. Δ=change. *p<0.05. Results expressed as mean ± SD.

*= participant (S6) was identified as the most influential point (Cook’s D >4) in the %-change in diaphragm MEP amplitudes, therefore, the data was not included in group analyses.*

**Table E1**: Measurements of PaCO2, PaO2 and mean arterial pressure (MAP) during baseline (BL), 30, 60, and 90 min post-AIHH in hApoE3 and hApoE4 rodent terminal neurophysiology experiment. Arterial PCO2 was maintained isocapnic (±1.5 mmHg) with baseline blood gas values. Baseline O2 levels (60% O2, balance N2 and CO2) were maintained for the duration of the experiments, except during the 15, 1-min hypoxic and hypercapnic episodes (Table E2; PaO2: 40-50 mmHg, 0.09 FiO2; PaCO2: 50-55 mmHg, +5-10 mmHg above baseline; 0.04 FiCO2; PETCO2: ≥ 48 mmHg).

| Groups |  | PaCO2, mmHg | | | | PaO2, mmHg | | | | MAP, mmHg | | | |
| --- | --- | --- | --- | --- | --- | --- | --- | --- | --- | --- | --- | --- | --- |
|  | n | BL | 30 min | 60 min | 90 min | BL | 30 min | 60 min | 90 min | BL | 30 min | 60 min | 90 min |
| **hApoE3** | 4 | 43.9 ± 1.5 | 44.0 ± 1.1 | 44.9 ± 0.9 | 44.4 ± 1.6 | 325 ± 17 | 220 ± 27 | 292 ± 5 | 277 ± 7 | 119 ± 4 | 119 ± 8 | 129 ± 2 | 131 ± 12 |
| **hApoE4** | 3 | 45.7 ± 1.2 | 45.6 ± 1.5 | 45.1 ± 2.2 | 46.2 ± 0.4 | 343 ± 11 | 284 ± 20 | 294 ± 17 | 278 ± 28 | 125 ± 2 | 118 ± 1 | 128 ± 9 | 130 ± 13 |

**Table E2**: Measurements of PaCO2, PaO2 and mean arterial pressure (MAP) during AIHH exposure. Blood gas measurements were taken at the 1st, 8th, and 15th hypoxic hypercapnia (HH) episode (HH1, HH8, and HH15, respectively).

| *Groups* |  | PaCO2, mmHg | | | PaO2, mmHg | | | MAP, mmHg | | |
| --- | --- | --- | --- | --- | --- | --- | --- | --- | --- | --- |
|  | n | HH1 | HH8 | HH15 | HH1 | HH8 | HH15 | HH1 | HH8 | HH15 |
| **hApoE3** | 4 | 49.0 ± 1.3 | 51.7 ± 1.1 | 52.0 ± 1.0 | 41.6 ± 0.8 | 45.1 ± 1.6 | 45.4 ± 1.5 | 93 ± 7 | 110 ± 12 | 111 ± 9 |
| **hApoE4** | 3 | 52.7 ± 1.4 | 53.6 ± 0.9 | 52.8 ± 0.9 | 42.5 ± 2.1 | 43.1 ± 2.2 | 46.4 ± 1.1 | 107 ± 13 | 101 ± 4 | 116 ± 14 |
